# Rpgrip1l controls ciliary gating by ensuring the proper amount of Cep290 at the vertebrate transition zone

**DOI:** 10.1101/2020.02.10.942300

**Authors:** Antonia Wiegering, Renate Dildrop, Christine Vesque, Sylvie Schneider-Maunoury, Christoph Gerhardt

## Abstract

A range of severe human diseases called ciliopathies are caused by the dysfunction of primary cilia. Primary cilia are cytoplasmic protrusions consisting of the basal body (BB), the axoneme and the transition zone (TZ). The BB is a modified mother centriole from which the axoneme, the microtubule-based ciliary scaffold, is formed. At the proximal end of the axoneme, the TZ functions as the ciliary gate governing ciliary protein entry and exit. Since ciliopathies often develop due to mutations in genes encoding proteins that localise to the TZ, the understanding of the mechanisms underlying TZ function is of eminent importance. Here, we show that the ciliopathy protein Rpgrip1l governs ciliary gating by ensuring the proper amount of Cep290 at the vertebrate TZ. Further, we identified the flavonoid eupatilin as a potential agent to tackle ciliopathies caused by mutations in *RPGRIP1L* as it rescues ciliary gating in the absence of Rpgrip1l.

## Introduction

The spatiotemporal regulation of cellular processes such as proliferation, apoptosis, migration, specification and differentiation depends on the cells’ ability to transduce signals from the environment into the cell’s interior. In nearly all mammalian cells, the primary cilium is dedicated to signal reception and transduction. Consequently, dysfunctional primary cilia result in severe often deadly human diseases, collectively called ciliopathies [1]. The current treatment of ciliopathies is restricted to symptomatic therapies and a curative medication against ciliopathies is missing [2]. In many cases, ciliopathies are caused by mutations in genes encoding TZ proteins [3, 4]. As the TZ functions as the ciliary gatekeeper governing ciliary protein import and export [4–12], a defective TZ can affect the proper formation of cilia and alter signalling pathways transduction [13–24]. Thus, the investigation of the molecular mechanisms underlying TZ function is an important prerequisite for the development of curative ciliopathy therapies.

In this study, we shed light on the role of Rpgrip1l in regulating the ciliary gating function of the TZ. Our previous investigations revealed that mutations in *RPGRIP1L* cause deadly ciliopathies [25], that Rpgrip1l localises to the vertebrate TZ [26] and that it is a decisive factor in vertebrate TZ assembly [27]. Our current work demonstrates that Rpgrip1l deficiency results in an altered ciliary protein composition and that Rpgrip1l governs ciliary gating by ensuring the proper amount of Cep290 at the vertebrate TZ. Further, we revealed that the flavonoid eupatilin rescues ciliary gating in the absence of Rpgrip1l. Consequently, eupatilin might represent a potential agent for the development of therapies against ciliopathies caused by mutations in *RPGRIP1L*.

## Results

### Rpgrip1l and Cep290 but not Nphp1, Nphp4 and Invs function as ciliary gatekeepers in mouse embryonic fibroblasts

Among the proteins allowed to cross the TZ are receptors and mediators of signalling pathways essential for proper development. Examples for such proteins are ADP Ribosylation Factor Like GTPase 13B (Arl13b), Somatostatin Receptor 3 (Sstr3), Smoothened (Smo), Polycystin 2 (Pkd2) or Adenylate Cyclase 3 (Ac3) which are often used as indicators to evaluate whether the gate function of the TZ is impaired [13, 14, 17, 19, 28–31]. In a former study, we showed that Rpgrip1l deficiency leads to a reduction of the ciliary Arl13b amount in all analysed mouse cells *in vitro* and *in vivo* [in mouse embryonic fibroblasts (MEFs), in mouse embryonic kidneys and in mouse limb buds][27]. However, we could not detect an alteration of the ciliary Smo amount in *Rpgrip1l*^−/−^ MEFs [26] raising the question whether the effect of Rpgrip1l is Arl13b-specific or whether it functions as a more general ciliary gatekeeper at the vertebrate TZ. In order to answer this question, we analysed the ciliary Sstr3 amount. It has been known for a long time that Sstr3 localises to cilia of neuronal cells [32]. More recently, it was shown that Sstr3 is also present in cilia of retinal pigment epithelial (RPE-1) cells [33]. We were now able to visualise endogenous Sstr3 in cilia of MEFs. Importantly, the amount of Sstr3 was decreased in *Rpgrip1l*^−/−^ MEFs (Figure 1A) demonstrating that Rpgrip1l exerts a TZ gatekeeper function. We next aimed to investigate how Rpgrip1l implements this function. Several proteins function as ciliary gatekeepers and it is an important task to unveil possible relationships between these gatekeeper proteins in order to understand the mechanisms which govern ciliary protein composition. In this context, Centrosome And Spindle Pole-Associated Protein 1 (Cspp1) was an interesting object of investigation since its truncation in humans results in a reduced ciliary amount of Arl13b and Ac3 [34] and since it interacts with Rpgrip1l [35]. Previously, we and others reported that Rpgrip1l ensures the proper amount of many proteins at the base of vertebrate cilia [19, 27] suggesting that it serves as a vital scaffold protein. Thus, we quantified the amount of Cspp1 at the ciliary base of *Rpgrip1l*^−/−^ MEFs but we could not detect any alteration (Figure S1A) indicating that Rpgrip1l exerts its ciliary gatekeeper function independently of Cspp1. Recently, a potential link between Rpgrip1l and the septin proteins was discussed [36]. Septins are known ciliary gatekeepers in vertebrates localising at the proximal end of the axoneme in MEFs [28]. We measured the ciliary amount of two different septins, Septin 2 (Sept2) and Septin 7 (Sept7), in *Rpgrip1l*^−/−^ MEFs. Neither Sept2 nor Sept7 was altered by the loss of Rpgrip1l (Figure S1B and C). We then turned to the TZ proteins Nephrocystin 4 (Nphp4), Nephrocystin 1 (Nphp1), Centrosomal Protein 290 (Cep290) and Inversin (Invs) whose amount at the vertebrate TZ is decreased in the absence of Rpgrip1l [27]. Previous studies in the invertebrates *Chlamydomonas reinhardtii* and/or *Caenorhabditis elegans* demonstrated that ciliary gating is regulated by the TZ proteins Nphp4, Nphp1, Cep290 and Invs [12, 17, 21, 23, 37] raising the possibility that Rpgrip1l might govern ciliary gating by controlling the amount of one of these proteins at the TZ. However, it is unknown whether these proteins function as ciliary gatekeepers in vertebrates and hence we quantified the ciliary amount of Arl13b and Sstr3 in *Nphp4*^−/−^, *Nphp1*^−/−^, *Cep290*^−/−^ and *Invs*^−/−^ mouse cells. The ciliary amount of both Arl13b and Sstr3 was unaffected in *Nphp4*^−/−^ MEFs (Figure 1B). To analyse TZ assembly, we had previously inactivated *Nphp1*, *Cep290* and *Invs* in NIH3T3 cells (immortalised MEFs) [27]. In the current study, we used these cells to investigate a possible involvement of these proteins in ciliary gating. To be able to perform comparative analyses in this context, we also inactivated Rpgrip1l in NIH3T3 cells (Figure S2A-C). As observed in MEFs [26], ciliary length was increased in *Rpgrip1l*^−/−^ NIH3T3 (Figure S2D). Moreover, the ciliary amount of Arl13b and Sstr3 was decreased in *Rpgrip1l*^−/−^ NIH3T3 (Figure 2A and B). In contrast to these cells, *Nphp1*^−/−^ and *Invs*^−/−^ NIH3T3 cells did not show an altered ciliary amount of Arl13b and Sstr3 (Figure 2A and B) indicating that the decreased amount of Nphp1 and Invs in *Rpgrip1l*^−/−^ MEFs is not the reason for the reduced ciliary Arl13b and Sstr3 amount. Importantly, Cep290 deficiency causes a decrease of the ciliary Arl13b and Sstr3 amount (Figure 2A and B) making it conceivable that the reduced amount of Cep290 might provoke the diminished amount of Arl13b and Sstr3 in the absence of Rpgrip1l.

**Figure 1:**
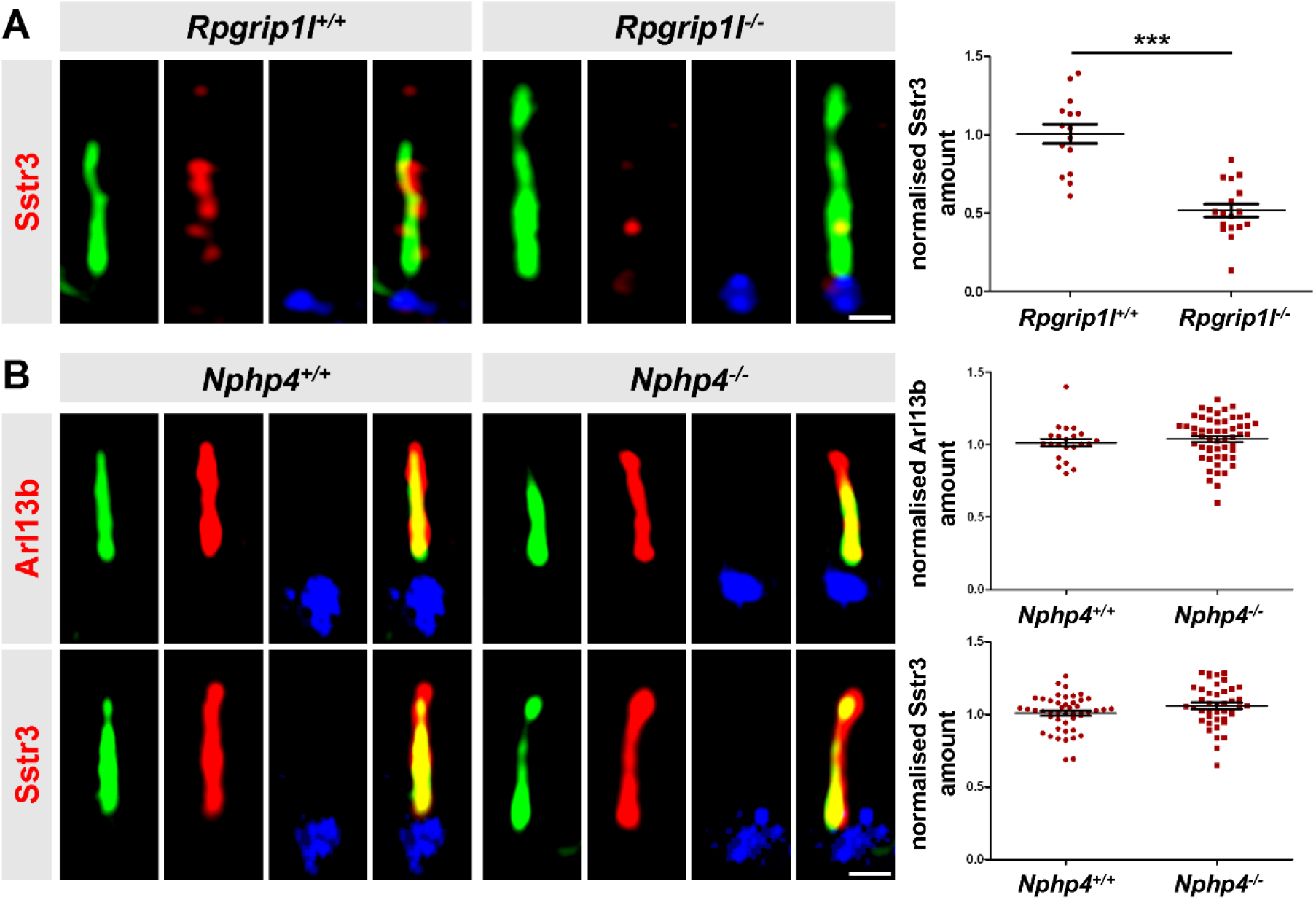
In mouse cells, ciliary gating is disturbed by the loss of Rpgrip1l but not by the loss of Nphp4. (A) Immunofluorescence on MEFs obtained from WT (n=5) and *Rpgrip1l*^−/−^ (n=5) embryos. (B) Immunofluorescence on MEFs obtained from WT (Arl13b: n=4; Sstr3: n=3) and *Nphp4*^−/−^ (Arl13b: n=4; Sstr3: n=3) embryos. (A, B) The ciliary axoneme is stained in green by acetylated α-tubulin, the basal body is stained in blue by γ-tubulin. The scale bars represent a length of 0.5 μm.

**Figure 2:**
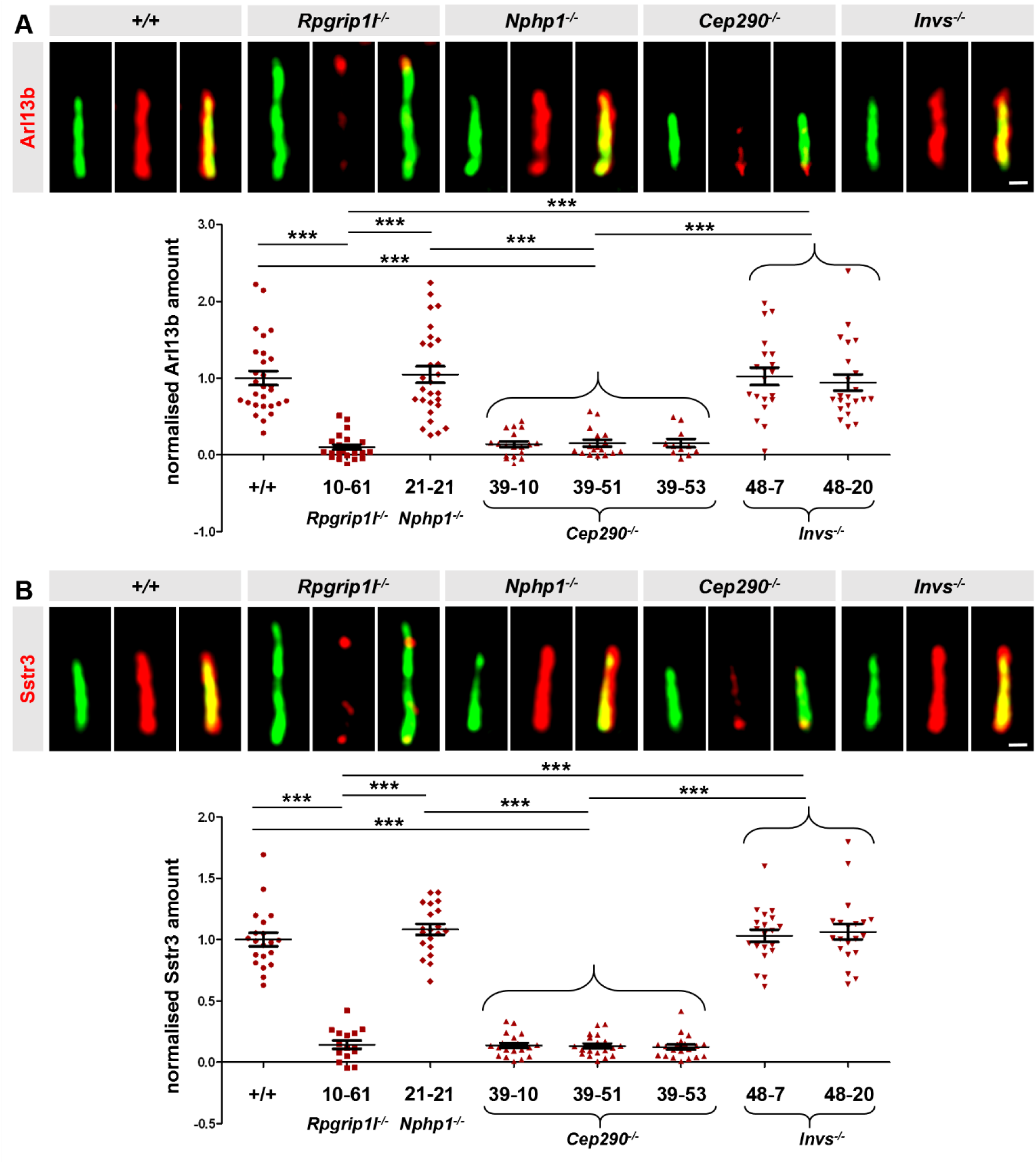
Loss of Cep290 but not of Nphp1 or Invs impairs ciliary gating in mouse cells. (A, B) Immunofluorescence on NIH3T3 cells. The ciliary axoneme is stained in green by acetylated α-tubulin. The scale bars represent a length of 0.5 μm.

### Restoration of the Cep290 amount at the TZ of *RPGRIP1L*^−/−^ HEK293 cells rescues the ciliary Arl13b amount

To test this hypothesis, we tried to enhance the amount of Cep290 at the TZ of *Rpgrip1l*^−/−^ MEFs and NIH3T3 cells by transfecting a plasmid that encodes a Flag-mCep290 fusion protein. Since the transfection failed in these cells, we transfected the plasmid into *RPGRIP1L*^−/−^ HEK293 cells. In a previous study, we revealed a reduced amount of Cep290 at the TZ of *RPGRIP1L*^−/−^ HEK293 cells [27]. The HEK293 cell line is an easily transfectable cell line [38] and, indeed, we detected the Flag-mCep290 fusion protein in *RPGRIP1L*^−/−^ HEK293 cells after transfection (Figure 3A). Remarkably, the fusion protein was located at the TZ (Figure 3B). In contrast to the Flag-mCep290 fusion protein, we could not detect a transfected Myc-mNphp1 fusion protein at the TZ of *RPGRIP1L*^−/−^ HEK293 cells (Figure S3A and B). Most likely, Rpgrip1l functions as a TZ scaffold for Nphp1 but not for Cep290.

**Figure 3:**
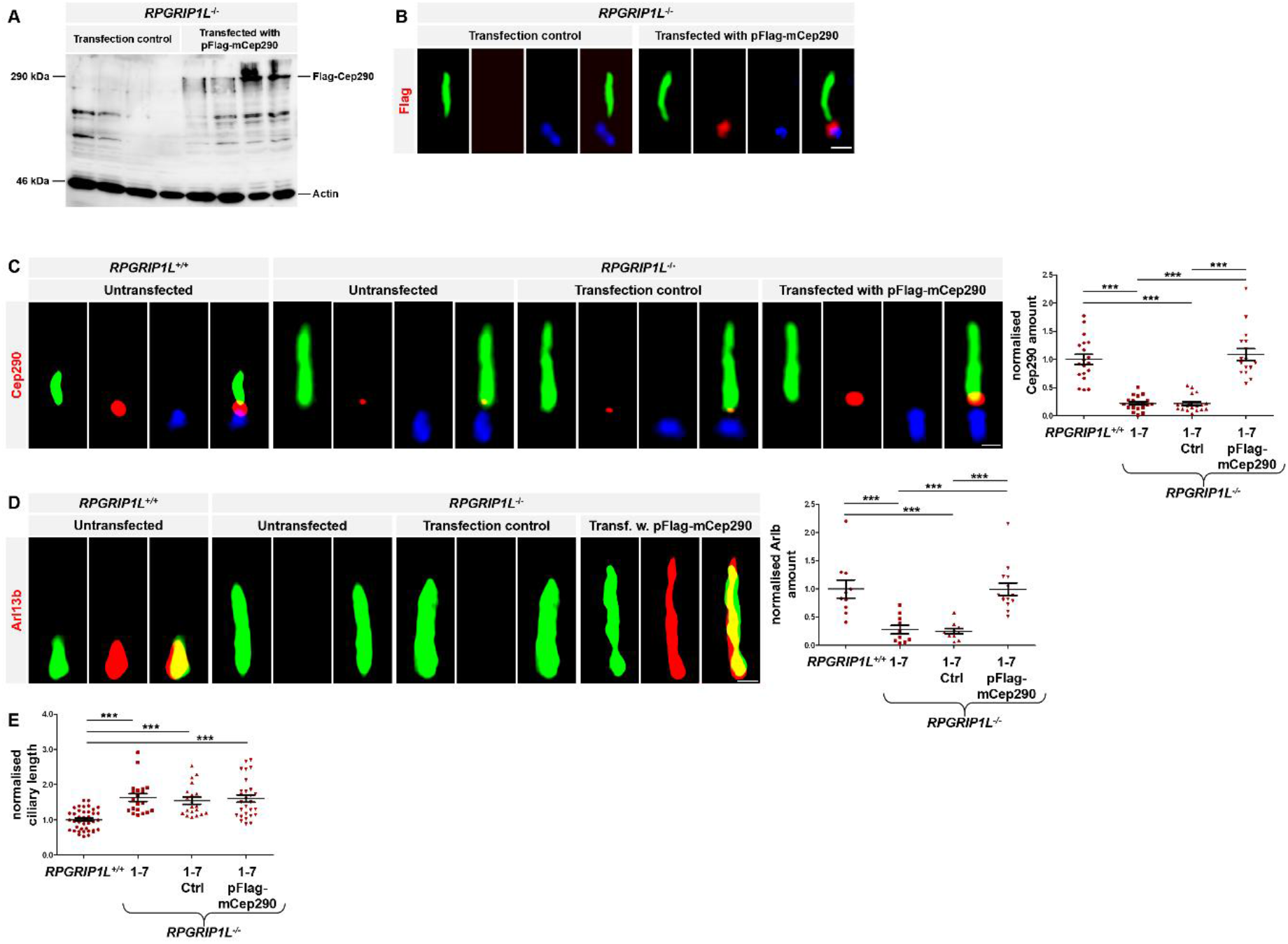
Rescue of the Cep290 amount at the TZ of *RPGRIP1L*^−/−^ HEK293 cells restores the ciliary Arl13b amount. (A) Western blot analysis with lysates obtained from *RPGRIP1L*^−/−^ HEK293 cells (clone 1-7). (B-D) Immunofluorescence on *RPGRIP1L*^+/+^ HEK293 cells and *RPGRIP1L*^−/−^ HEK293 cells (clone 1-7). The ciliary axoneme is stained in green by acetylated α-tubulin, the basal body is stained in blue by γ-tubulin (B, C). The scale bar represents a length of 1 μm (B) or 0.5 μm (C, D). (E) Ciliary length quantification.

The transfection of the plasmid encoding the Flag-mCep290 fusion protein into *RPGRIP1L*^−/−^ HEK293 cells, rescued the amount of Cep290 at the TZ (Figure 3C). Importantly, this rescue restored the ciliary amount of Arl13b in *RPGRIP1L*^−/−^ HEK293 cells (Figure 3D) indicating that the decreased amount of Cep290 in *RPGRIP1L*^−/−^ HEK293 cells causes an impaired ciliary gating. Cilia of HEK293 cells are elongated by the loss of Rpgrip1l [27]. Interestingly, the increased cilia length in the absence of Rpgrip1l was not rescued by the expression of the Flag-mCep290 fusion protein (Figure 3E).

### Eupatilin treatment rescues ciliary gating in Rpgrip1l-negative mouse embryonic fibroblasts

HEK293 cells lack Sstr3, preventing the analysis of its localisation in the absence of Rpgrip1l and its rescue by Cep290 overexpression [39]. For this reason, we performed another approach by using the flavonoid eupatilin. A recent report showed that eupatilin rescues the ciliary gating in *CEP290*^−/−^ human cells ^6^. The treatment of *Rpgrip1l*^−/−^ NIH3T3 cells with eupatilin restored the amount of both Arl13b and Sstr3 (Figure 4A and B) indicating that Rpgrip1l controls ciliary gating via ensuring the proper amount of Cep290 at the vertebrate TZ. The enhanced cilia length in *Rpgrip1l*^−/−^ NIH3T3 cells was not rescued by eupatilin (Figure 4C).

**Figure 4:**
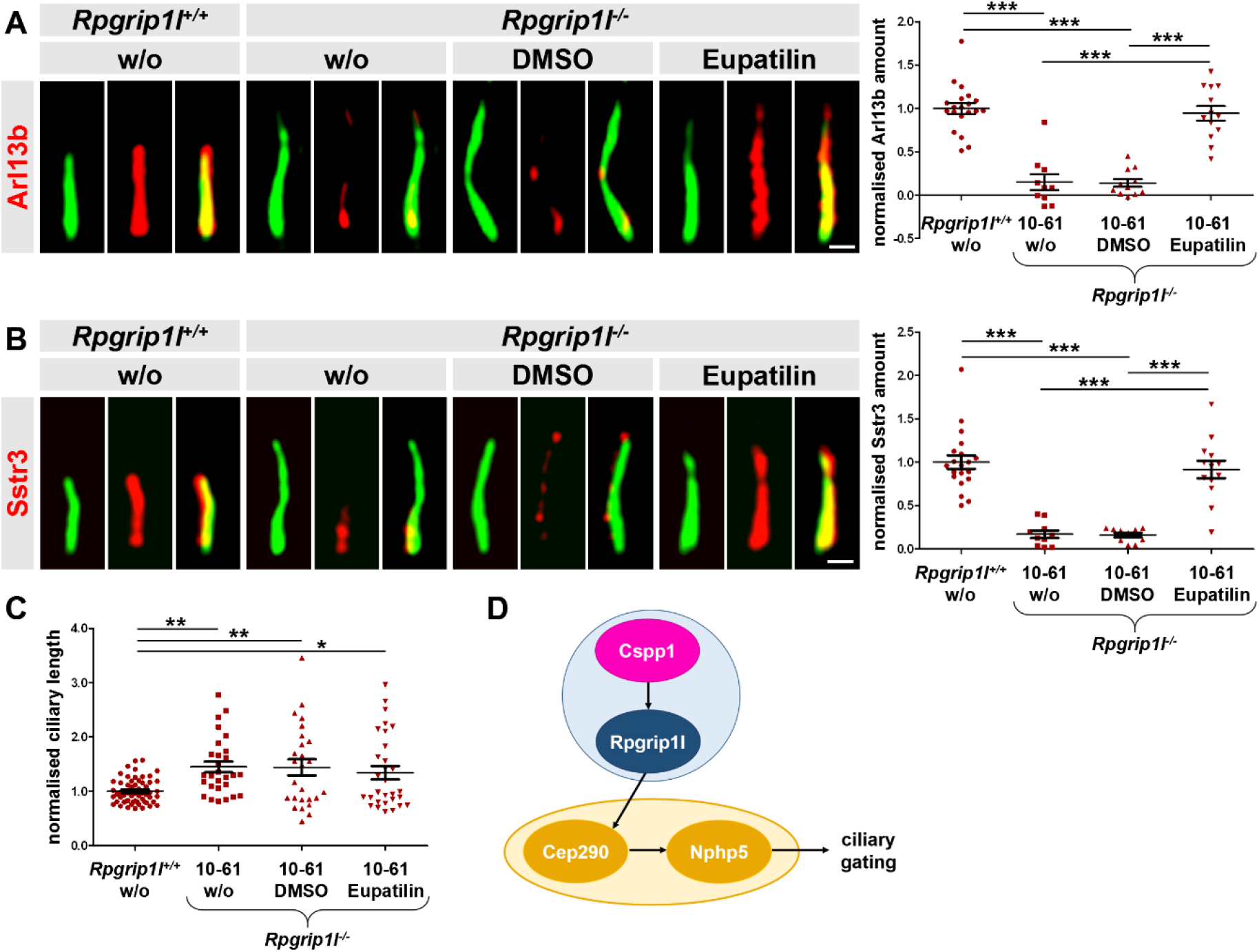
Eupatilin treatment rescues ciliary gating in the absence of Rpgrip1l. (A, B) Immunofluorescence on *RpgrIP1L*^+/+^ and *Rpgrip1l*^−/−^ NIH3T3 cells. The ciliary axoneme is stained in green by acetylated α-tubulin. The scale bars represent a length of 0.5 μm. (C) Ciliary length quantification. (D) Model of ciliary gating hierarchy. Interaction partners are depicted in circles.

## Discussion

Primary cilia mediate numerous signalling pathways thereby ensuring proper development and homeostasis. In this context, the intraciliary concentration of proteins involved in these signalling pathways is of enormous importance. Consequently, ciliary import and export and hence ciliary protein composition has to be tightly controlled. This control is implemented by the TZ. Since mutations in genes encoding TZ proteins result in ciliopathies [1, 3, 4], current cilia research aims to uncover mechanisms underlying TZ assembly and function. However, little is known about these mechanisms in vertebrates. Recently, we described Rpgrip1l as a decisive factor in vertebrate TZ assembly [27]. In this context, Rpgrip1l deficiency leads to a reduced amount of Cep290, Nphp1, Nphp4 and Invs at the TZ [27]. Rpgrip1l, Cep290, Nphp1, Nphp4 and Invs were previously shown to govern ciliary gating in *Chlamydomonas reinhardtii* and/or *Caenorhabditis elegans* [12, 17, 21, 23, 37, 40]. However, loss of Nphp1, Nphp4 and Invs did not alter the ciliary amount of Arl13b or Sstr3 in MEFs and NIH3T3 cells (Figure 1B and Figure 2) indicating that they are not involved in gating these proteins in vertebrate primary cilia. Remarkably, several reports point to cell type-specific functions of some TZ proteins [14, 16, 27, 41, 42] making a potential regulation of ciliary gating by Nphp1, Nphp4 and Invs in other vertebrate cell types conceivable. *Rpgrip1l*^−/−^ and *Cep290*^−/−^ mice have a much more severe phenotype than *Nphp1*^−/−^, *Nphp4*^−/−^ and *Invs*^−/−^ mice [20, 25, 27, 42–59]. Moreover, mutations in *RPGRIP1L* and *CEP290* result in more severe human ciliopathies than mutations in *NPHP1*, *NPHP4* or *INVS* [60–63]. On the one hand, these differences might reflect that Rpgrip1l and Cep290 function as ciliary gatekeepers in vertebrates while Nphp1, Nphp4 and Invs do not. On the other hand, these differences might be based on the fact that Rpgrip1l and Cep290 exert additional functions in the cytoplasm, e.g. the regulation of protein degradation systems [26, 64] or the organisation of the cytoplasmic microtubule network [65].

Formerly, we demonstrated that Rpgrip1l deficiency does not affect the overall cellular amount of Cep290 but its localisation at the vertebrate TZ [27]. There is a perennial debate about the function(s) of Rpgrip1l. Does it predominantly serve as a structural TZ anchor or scaffold protein and interacts with other proteins thereby ensuring their localisation and proper amount at the TZ or does it control the TZ localisation and amount of proteins by exerting additional functions e.g. regulating protein degradation systems, functioning as a TZ assembly factor, establishing a ciliary zone of exclusion (CIZE) that excludes signal transduction proteins etc. [15, 19, 23, 26, 27, 64, 66–68]? In this study, the Flag-mCep290 protein was able to localise to the TZ in the absence of Rpgrip1l (Figure 3B) indicating that Rpgrip1l does not function as a structural scaffold for the TZ presence of Cep290. In line with this assumption, it was not shown yet that Rpgrip1l interacts with Cep290. To stress this point, we also transfected a plasmid encoding a Myc-mNphp1 fusion protein into *RPGRIP1L*^−/−^ HEK293 cells. It was reported before that Nphp1 interacts with Rpgrip1l [69]. Since Myc-mNphp1 was not present at the TZ in the absence of Rpgrip1l (Figure S3B), we suggest that Rpgrip1l functions as structural anchor for Nphp1 but not for Cep290. Dissecting the mechanism by which Rpgrip1l regulates the amount of Cep290 at the vertebrate TZ will be a thrilling future project.

Here, we show that Rpgrip1l controls ciliary gating via ensuring the proper amount of Cep290 at the TZ (Figure 3D). Cep290 governs ciliary protein composition by interacting with Nphp5 [70]. Consequently, Rpgrip1l, Cep290 and Nphp5 can be defined as ciliary gatekeepers. Cep290 binds to Nphp5 thereby preventing the binding of calmodulin to Nphp5 and promoting the recruitment of Nphp5 to the TZ [71]. In the absence of Cep290, Nphp5 is not present at the TZ. However, the treatment of *CEP290*^−/−^ cells with eupatilin rescues the TZ amount of Nphp5 and ciliary gating since eupatilin inhibits the binding between Nphp5 and calmodulin [71]. In the current study, eupatilin treatment rescued ciliary gating in the *Rpgrip1l* deficient state (Figure 4A and B). For this reason, we hypothesise that Rpgrip1l controls the proper TZ amount of Nphp5 by its effect on Cep290. In turn, Nphp5 ensures ciliary gating (Figure 4D). Thus, we propose eupatilin as a potential agent for the treatment of ciliopathies caused by mutations in *RPGRIP1L*. Interestingly, Rpgrip1l was not required for the ciliary localisation of Cspp1 Figure S1A) but the proper amount of Rpgrip1l at the TZ depends on Cspp1 [35]. Moreover, mutations in *CSPP1* disturb ciliary protein composition (e.g. reduced ciliary Arl13b amount) [34] and cause Joubert syndrome and Meckel syndrome [72]. Based on these facts, we place Cspp1 at the top of the “ciliary gating hierarchy” involving Rpgrip1l, Cep290 and Nphp5 (Figure 4D).

Interestingly, Garcia-Gonzalo et al. revealed that the loss of the TZ gatekeeper protein Tmem67 diminishes the ciliary amount of Arl13b and Ac3 but the ciliary amount of Smo remains normal [14] demonstrating the existence of a specificity between the gatekeeper proteins and the proteins which are allowed to cross the TZ. In *Rpgrip1l*^−/−^ MEFs, the ciliary amount of Smo is also unaltered [26]. The lack of Cep290 and Nphp5 in RPE-1 cells results in a reduced ciliary amount of Smo [70]. Since the amount of Cep290 is reduced in *Rpgrip1l*^−/−^ MEFs [27], the expectation would be that the amount of Smo was decreased in these MEFs. However, these findings have been made in different cell types making it possible that the ciliary gating function of these proteins might be cell type-specific. The analysis of this hypothesis is an exciting subject of future studies which would shed further light on the ciliary gating function of the TZ.

Many ciliopathies can be attributed to mutations in genes encoding TZ proteins [3, 4]. For this reason, the assembly and function of the TZ is a hot topic in biomedical research. A lot of proteins participate in TZ assembly and/or function as ciliary gatekeepers at the TZ [12–14, 16, 17, 19, 23, 24, 27, 29, 30, 33, 34, 37, 41, 69, 70, 73–90]. However, the relationships between these proteins and hence the mechanisms underlying ciliary gating at the TZ remain largely elusive. Recently, we showed that Rpgrip1l represents a central factor in vertebrate TZ assembly [27]. Our current study reveals that Rpgrip1l also regulates ciliary gating by ensuring the proper amount of Cep290 at the vertebrate TZ. Combining our results with previous findings, we suggest a protein hierarchy regulating ciliary gating in which Cspp1, Rpgrip1l, Cep290 and Nphp5 are involved. Our work is an important piece of a puzzle depicting this fundamental ciliary process. The completion of this puzzle will be one of the most important tasks of cilia research in the next few years.

## Materials and Methods

### Cell lines

We used two different cell lines in this study. NIH3T3 cells (#ACC59) and HEK293 cells (#ACC35), both purchased by the German Collection of Microorganisms and Cell Cultures GmbH (DSMZ). Cells were grown in DMEM supplemented with 10% fetal calf serum (FCS), 1/100 (v/v) L-glutamine (Gibco), 1/100 (v/v) sodium pyruvate (Gibco), 1/100 (v/v) non-essential amino acids (Gibco) and 1/100 (v/v) pen/strep (Gibco) at 37 °C and 5% CO_2_. The following clones were used: *Rpgrip1l*^−/−^ NIH3T3 cells (clone 10-61), *Cep290*^−/−^ NIH3T3 cells (clones 39-10, 39-51, 39-53) **[27]**, *Nphp1*^−/−^ NIH3T3 cells (clone 21-21) **[27]**, *Invs*^−/−^ NIH3T3 cells (clones 48-7, 48-20) **[27]** and *RPGRIP1L*^−/−^ HEK293 cells (clone 1-7) **[27]**.

### Primary Cell Culture

We isolated MEFs from single mouse embryos (male and female) at embryonic stage (E) 12.5 after standard procedures. MEFs were grown in DMEM supplemented with 10% fetal calf serum (FCS), 1/100 (v/v) L-glutamine (Gibco), 1/100 (v/v) sodium pyruvate (Gibco), 1/100 (v/v) non-essential amino acids (Gibco) and 1/100 (v/v) pen/strep (Gibco) at 37 °C and 5% CO_2_. The following mutant mice were used: *Rpgrip1l*-mutant mice on a C3H-background **[20]** and *Nphp4*-mutant mice on a C57BL/6J-background **[27]**.

### Method Details

#### Cell culture, transfection and drug treatment

Ciliogenesis in confluent grown MEFs and NIH3T3 cells was induced by serum-starvation (0.5% FCS) for at least 24 hours. For DNA transfection, Lipofectamin 3000 (Invitrogen) was used following the manufactures guidelines. NIH3T3 cells were treated with 20 μM eupatilin (#SML1689; Sigma-Aldrich) or DMSO as a solvent control for 24 h.

#### Antibodies and Plasmids

Cells were immmunolabeled with primary antibodies targeting actin (#A2066; Sigma-Aldrich), Arl13b (#17711-1-AP; Proteintech), Cep290 (#ab84870; Abcam), Cspp1 (#11931-1-AP; Proteintech), Flag (#F7425; Sigma-Aldrich), Myc (#sc-789; Santa Cruz Biotechnology, Inc.), Sept2 (#11397-1-AP; Proteintech), Sept7 (#13818-1-AP; Proteintech), Sstr3 (#20696-1-AP; Proteintech and #PA3-207; Pierce Biotechnology), acetylated α-tubulin (#sc-23950; Santa Cruz Biotechnology, Inc.), and γ-tubulin (#sc-7396; Santa Cruz Biotechnology, Inc.). The generation of the polyclonal antibody against Rpgrip1l was described formerly **[20]**.

The following plasmids were used: pMyc-mNphp1 (kindly provided by Sophie Saunier) and pFlag-mCep290 (#27381; Addgene). pMyc-mNphp1 encodes for the murine full-length Nphp1 protein fused to a myc-tag (vector: CMV) and pFlag-mCep290 encodes for the murine full-length Cep290 protein fused to a flag-tag (vector: CMV2).

#### CRISPR/Cas9-mediated Gene Inactivation

Inactivation of mouse *Rpgrip1l* in NIH3T3 cells was performed as previously described **[27]**. We choose a target site which is located in exon3 of the gene (Figure S2). After inactivation and single-cell cloning, 8 clones, which upon RFLP analysis appeared to have lost the diagnostic *Eag*I recognition sequence, were further analysed. To establish the genotype, individual alleles were cloned and sequenced (Figure S2).

Sequences of the target site and primer pairs used to amplify the targeted region are as follows:

Rpgrip1l ‒T3: CTCGAGTTAACACCGGCCGCCGG
Rpgrip1l-T3b-for: GAATGGCCACCAAGTTAATACGGCTAG
Rpgrip1l-T3b-rev: CTTCAGGATCTGACAGAGAGCAAGCCTC

#### Off-target Analyses

Off-target analyses were performed by RFLP analyses as previously described **[27]**. For the on-target Rpgrip1l-T3 there exist no off-targets carrying either one, two or three mismatches. From the remaining 4-mismatch off-targets we tested the top-3 ranking sites **[91]** (crispr.mit.edu/) on the DNAs from the same 8-set-clones analysed for targeting of the on-target. In summary, we did not detect any mutations (data not shown).

Sequences of the off-target sites and primer pairs used for amplification are as follows:

T3 offtarget-1: CACGAGTCAGCACCGGCCACTGG
T3 off-1 for: CTGTCAGGTTTCCCAGTGTGCAG
T3 off-1 rev: CTCTCAGCTCCTTTTAGGTCTCCAG
T3 offtarget-2: CTCTACTGAACAACGGCCGCAGG
T3 off-2 for: ATCCAGCCAAACCCTGCCTGTTC
T3 off-2 rev: GGTTTGTCTCTGTCCTGACATGTCAC
T3 offtarget-3: ATCCAGTTGACACCGGCCTCTGG
T3 off-3 for: GTCTCCTTCAGACCCACTGAAGTG
T3 off-3 rev: GTCCCAGGAAGCCAGGCTGTTG

The following restriction enzymes were used: *Bsl*I (T3 offtarget-1), *Eag*I (T3 offtarget-2), *Bsl*I (T3 offtarget-1).

#### Image Processing

Image acquisition and data analysis were carried out at room temperature using a Zeiss Imager.A2 microscope, 100x, NA 1.46 oil immersion objective lens (Carl Zeiss AG), a monochrome charge-coupled device camera (AxioCam MRm, Carl Zeiss AG), and the AxioVision Rel. 4.8 software package (Carl Zeiss AG). Appropriate anti-mouse, anti-rabbit and anti-goat Alexa405, Cy3, and Alexa488 antibodies were used as fluorochromes.

#### Immunofluorescence

For immunofluorescence on MEFs, NIH3T3 cells and HEK293 cells, cells were plated on coverslips until confluency. MEFs and NIH3T3 cells were serum-starved for at least 24 hours. Cells were fixed with 4% PFA (for stainings with the antibodies to Cep290, Rpgrip1l, Flag, Myc, Sept7 and Arl13b) or methanol (for stainings with the antibodies to Cspp1, Sept2, Sstr3). Fixed cells were rinsed three times with PBS, followed by a permeabilisation step with PBS/0.5% Triton-X-100 for 10 minutes. The samples were rinsed three times with PBS. Samples were incubated for at least 10 minutes at room temperature in PBST (PBS/0.1% Triton-X-100) containing 10% donkey serum. Diluted primary antibodies (in blocking solution) were incubated overnight at 4 °C. After three washing steps with PBST, incubation with fluorescent secondary antibody (diluted in blocking solution) was performed at room temperature for 1 hour followed by several washing steps and subsequent embedding with Mowiol.

#### Western Blotting

Whole-cell lysates were obtained by lysis with radioimmunoprecipitation buffer (150 mM sodium chloride, 50 mM Tris-HCl, pH 7.4, 0.1% sodium deoxycholate, and 1 mM EDTA). Protein content was measured by the Bradford method, and samples were normalized. 20 mg of total protein was separated by SDS-PAGE on polyacrylamide gels (10%) and transferred to a polyvinylidene fluoride membrane (Bio-Rad Laboratories, Inc.). The membrane was probed with antibodies against Flag (#F7425; Sigma-Aldrich) and Myc (#sc-789; Santa Cruz Biotechnology, Inc.). Anti-actin (#A2066; Sigma-Aldrich) antibody was used as loading control. Proteins were detected with secondary antibodies conjugated to horseradish peroxidase (RPN4201 and RPN4301) and the ECL detection kit (both GE Healthcare). Visualization of protein bands was realized by LAS-4000 mini (Fujifilm). Band intensities were measured by using ImageJ (National Institutes of Health).

### Quantification and Statistical Analysis

#### Quantification

Quantifications of ciliary protein staining and protein bands intensity were quantified using ImageJ (National Institutes of Health). Intensity measurement of proteins based on immunofluorescence staining was performed as described before **[14, 24, 26, 27, 29, 64, 92]**. The ciliary length has to be taken into account while quantifying the ciliary amount of Arl13b and Sstr3 in different genotypes. Therefore we used the area marked by acetylated α-tubulin as a reference and quantified the average pixel intensity of the Arl13 and Sstr3 staining. For all other ciliary protein intensities (Cep290, Flag, Cspp1, Myc, Rpgrip1l, Sept2 and Sept7), we selected the region labelled by γ-tubulin (for BB proteins) or the area in-between the γ-tubulin staining and the proximal part of the acetylated α-tubulin staining and measured the total pixel intensity. To exclude unspecific staining from the measurements, we subtracted the mean value of the average pixel intensity (in the case of Arl13b and Sstr3) or of the total pixel intensity (all other ciliary proteins) of three neighboring regions free from specific staining.

#### Statistical Analysis

Data are presented as mean ± standard error of mean (SEM). Two-tailed *t-*test with Welch’s correction was performed for all data in which two datasets were compared. Analysis of variance (ANOVA) and Tukey honest significance difference (HSD) tests were used for all data in which more than two datasets were compared. A P-value <0.05 was considered to be statistically significant (one asterisk), a P-value <0.01 was defined as statistically very significant (two asterisks) and a P-value <0.001 was noted as statistically high significant (three asterisks). All statistical data analysis and graph illustrations were performed by using Graphpad Prism (Graphpad Software Inc). Sample sizes are indicated in the figure legends.

## Acknowledgments

The authors thank Matias Zurbriggen and Leonie-Alexa Koch for their generous help to enable the continuation of the study. Moreover, we are grateful to Sophie Saunier for providing the Myc-mNphp1 construct. This work was funded by the Fondation ARC pour la Recherche sur le Cancer (Project ARC PJA 20171206591 to S.S.M.) and the Fondation pour la Recherche Médicale (Equipe FRM EQU201903007943 to S.S.M.).

## Author contributions

Conceptualisation, C.G.; Methodology, R.D.; Validation, A.W., R.D. and C.G.; Formal Analysis, A.W.; Investigation, A.W., R.D. and C.G.; Resources, C.V. and S.S.M.; Writing – Original Draft, C.G.; Visualisation, A.W.; Supervision, C.G.; Project Administration, C.G.; Funding Acquisition, S.S.M.

