## Supporting Information File for "Rpgrip1l controls ciliary gating by ensuring the proper amount of Cep290 at the vertebrate transition zone"

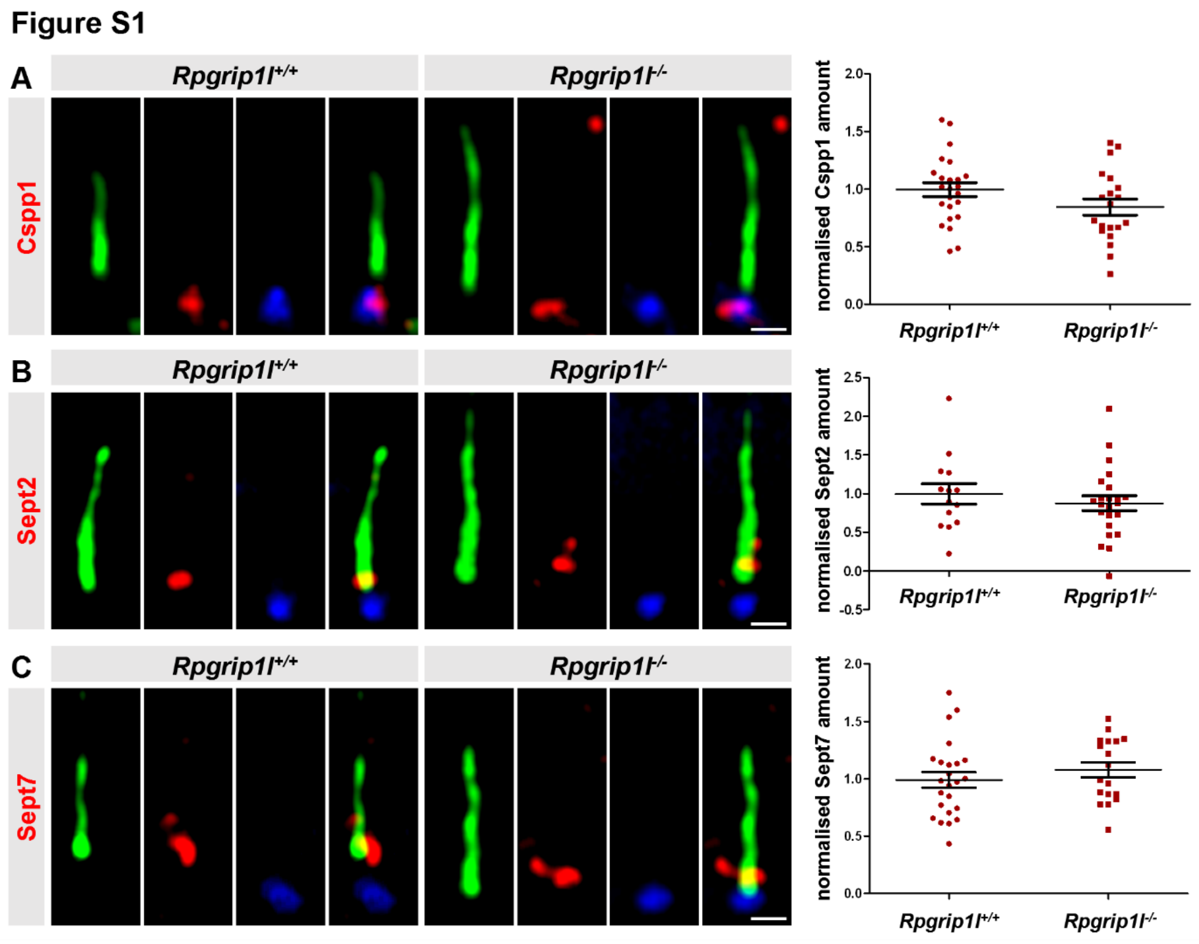


**Figure S1: The ciliary amount of Cspp1, Sept2 and Sept7 does not depend on Rpgrip1l.** (A-C) Immunofluorescence on MEFs obtained from WT (n=4) and *Rpgrip1l*$^{-/-}$ (n=4) embryos at embryonic stage E12.5. The ciliary axoneme is stained in green by acetylated α-tubulin, the basal body is stained in blue by γ-tubulin. The scale bars represent a length of 0.5 µm. The amounts of Cspp1, Sept2 and Sept7 are unaltered in cilia of *Rpgrip1l*$^{-/-}$ MEFs. Data are shown as mean ± s.e.m. Asterisks denote statistical significance according to unpaired *t*-tests with Welch's correction (A: $t$ (37) = 1.649, P < 0.1072; B: $t$ (26) = 1.088, P< 0.2867; C: $t$ (39) = 0.926, P < 0.3601).

#### Figure S2

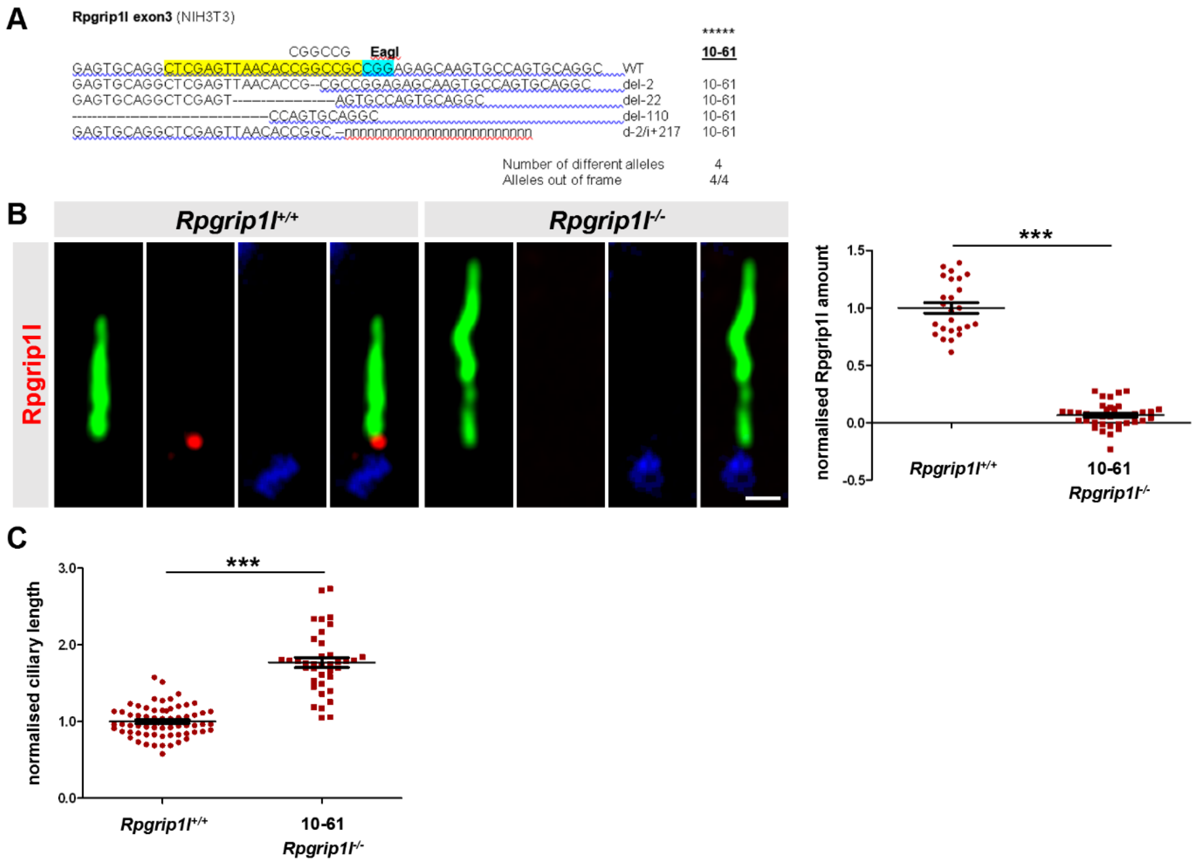


**Figure S2: Confirmation of the inactivation of *Rpgrip1l* in *Rpgrip1l*$^{-/-}$ NIH3T3 cells.**

(A) Genotype analysis of targeted *Rpgrip1l* on-target alleles. Sequences of CRISPR/Cas9 mutated alleles are compared to the WT sequence (on the left side), corresponding NIH clone 10-61 is depicted on the right side. The sequences of the 20 nt guide and the PAM within the WT sequence are coloured in yellow and cyan, respectively. The recognition sequence of *Eag*I (used in RFLP analyses to screen for mutated clones) is shown on top of the WT sequence. Within the sequences of targeted alleles, deletions are indicated by dashes, insertions of a single base pair are typed bold, and larger insertions are displayed as runs of "n". The karyotype analysis of NIH3T3 revealed, that *Rpgrip1l* is present in 4 copies within this cell line [93]. In accordance with this, we were able to detect four different alleles in clone 10-61 appeared to carry only alleles with out-of-frame mutations. (B) Immunofluorescence on *Rpgrip1l*$^{+/+}$ and *Rpgrip1l*$^{-/-}$ (clone 10.61) NIH3T3 cells. The ciliary axoneme is stained in green by acetylated α-tubulin, the basal body is stained in blue by γ-tubulin. The scale bars represent a length of 0.5 µm. Rpgrip1l is not detectable in cilia of *Rpgrip1l*$^{-/-}$ NIH3T3 cells (clone 10.61). Data are shown as mean ± s.e.m. Asterisks denote statistical significance according to an unpaired *t*-test with Welch's correction (***P < 0.001) ($t$ (30) = 18.72, P < 0.0001). (C) Ciliary length measurement. Data are shown as mean ± s.e.m. Asterisks denote statistical significance according to an unpaired *t*-test with Welch´s correction (***P < 0.001) ($t$ (47) = 11.06, P < 0.0001).

### Figure S3

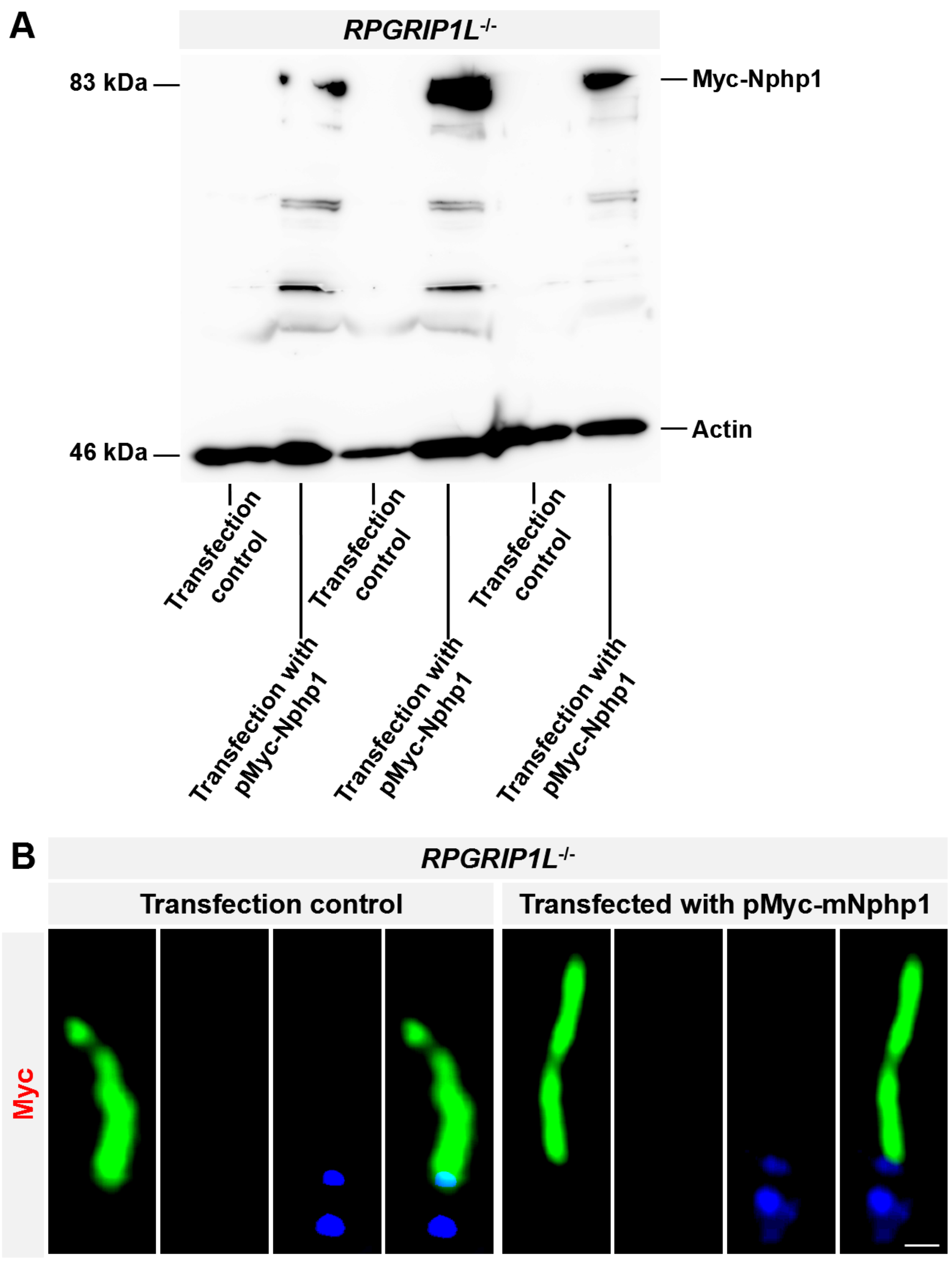


**Figure S3: The localisation of Nphp1 at the TZ depends on Rpgrip1l.**

(A) Western blot analysis with lysates obtained from *RPGRIP1L*$^{-/-}$ HEK293 cells (clone 1-7). Cells were transfected with a plasmid encoding a Nphp1 (full-length)-Myc fusion protein. Actin serves as loading control. (B) Immunfluorescence on *RPGRIP1L*$^{-/-}$ HEK293 cells (clone 1-7). The ciliary axoneme is stained in green by acetylated α-tubulin, the basal body is stained in blue by γ-tubulin. Myc is shown in red. The Nphp1-Myc fusion protein is not detectable at the TZ. The scale bar represents a length of 0.5 µm.
